# A method to enrich and purify centromeric DNA from human cells

**DOI:** 10.1101/2021.09.24.461328

**Authors:** Riccardo Gamba, Giulia Mazzucco, Therese Wilhelm, Florian Chardon, Leonid Velikovsky, Julien Picotto, Ylli Doksani, Daniele Fachinetti

## Abstract

Centromeres are key elements for chromosome segregation. Canonical centromeres are built over long-stretches of tandem repetitive arrays. Despite being quite abundant compared to other loci, centromere sequences overall still represent only 2 to 5% of the human genome, therefore studying their genetic and epigenetic features is a major challenge. Furthermore, sequencing of centromeric regions requires high coverage to fully analyze length and sequence variations, which can be extremely costly. To bypass these issues, we have developed a technique based on selective restriction digestion and size fractionation to enrich for centromeric DNA from human cells. Combining enzymes capable of cutting at high frequency throughout the genome, except within most human centromeres, with size-selection of >20 kb fragments resulted in over 25-fold enrichment in centromeric DNA. Sequencing of the enriched fractions revealed that up to 60% of the enriched material is made of centromeric DNA. This approach has great potential for making sequencing of centromeric DNA more affordable and efficient and for single DNA molecule studies.

## Introduction

Centromeres are the chromosomal site for assembly of the kinetochore, the fundamental complex necessary for proper chromosome segregation in both meiosis and mitosis (Balzano & Giunta, 2020; Das *et al*, 2020). In humans they are composed of highly repetitive arrays of alpha satellite DNA (α-sat) that stretches over megabase-long regions (Sullivan & Sullivan, 2020). α-sat DNA is organized in a head-to-tail tandem repeats of single AT-rich 171 bp monomers that can form highly homogeneous Higher Order Repeat (HOR) units of different length and composition among different chromosomes. These HORs are typically flanked by monomeric diverged alpha satellite repeats, and different HOR arrays on the same centromere can be separated by other repeat families (Altemose *et al*, 2021a; Logsdon *et al*, 2021; Miga *et al*, 2020; Nurk *et al*, 2021).

Centromeric DNA and its DNA binding protein CENP-B have been recently implicated in centromere stability or function (Gamba & Fachinetti, 2020; Ohzeki *et al*, 2020; Lampson & Black, 2017; Ali-Ahmad & Sekulić, 2020). Yet, the repetitive nature of these loci has hindered their detailed molecular characterization. The use of novel, long-read sequencing approaches and the development of new computational methods has recently allowed a breakthrough in the dissection of the sequence of these long repetitive regions. This is exemplified by the recent release of a whole uninterrupted telomere to telomere (T2T) sequence of a human genome (from a hydatidiform mole derived cell line, CHM13-TERT (Altemose *et al*, 2021a; Logsdon *et al*, 2021; Miga *et al*, 2020; Nurk *et al*, 2021). These progresses in DNA sequence mapping open a new era in the genomic studies of centromeres. Nevertheless, probing centromeric DNA still poses some difficulties, also considering that centromeric repeats can vary across individuals and between homologous chromosomes.

A big limitation in the study of centromeric DNA is that there are no widely-established and efficient methods to select centromeric regions and isolate them from the rest of the genome. Therefore, investigation of the centromeric sequence requires whole genome sequencing (WGS), a very inefficient and costly approach as only 2-5% of the human genome is composed by centromeric DNA (Aldrup-MacDonald & Sullivan, 2014; Altemose *et al*, 2021a). Further, single molecule studies on centromeric DNA aimed to study their replication and structure are either strongly limited by the usage of fluorescent probes to identify centromere DNA or totally lacking.

Use of immuno-precipitation methods relying on the presence of centromeric proteins can only isolate a sub-portion of the whole centromeric array. According to recent estimates, CENP-A, the histone H3 variant enriched at centromeric regions (McKinley *et al*, 2015), spans a region of approximately 0.2 to 0.5 Mb per centromere, totaling to ∼7.8Mb, less than 10% of the total alpha satellite content in the genome (Altemose *et al*, 2021a). Also, immuno-precipitation methods do not provide long, uninterrupted DNA fragments that are necessary to unravel the centromere sequence and structure.

Another approach to enrich for a target sequence is based on restriction enzymes and relies on the digestion of the rest of the genome while maintaining the regions of interest largely intact. This rationale is applied for the purification of telomeric repeats, which lack canonical restriction sites (de Lange *et al*, 1990; Mender & Shay, 2015; Griffith *et al*, 1999). More recently, a two-step procedure has been developed for the study of telomere structure by electron microscopy (EM) (Mazzucco *et al*, 2020). While a similar restriction-based approach was developed in the pre-genomic era to isolate mouse (peri)centromeres (Lica & Hamkalo, 1983), an analogous technique for the study of human centromeres is currently missing.

In this manuscript, we present the development of a restriction digestion-based method to enrich for centromeric repeats that allows isolation of high molecular weight (HMW), long fragments of centromeric DNA suitable for long-read sequencing (**Figure 1A**). Our method drastically increases the efficiency of sequencing centromeric DNA compared to whole genome sequencing. Furthermore, we demonstrate that this method allows direct visualization of long centromeric fragments in fluorescence microscopy, with possible applications for single-molecule analysis of centromeric DNA.

**Figure 1.**
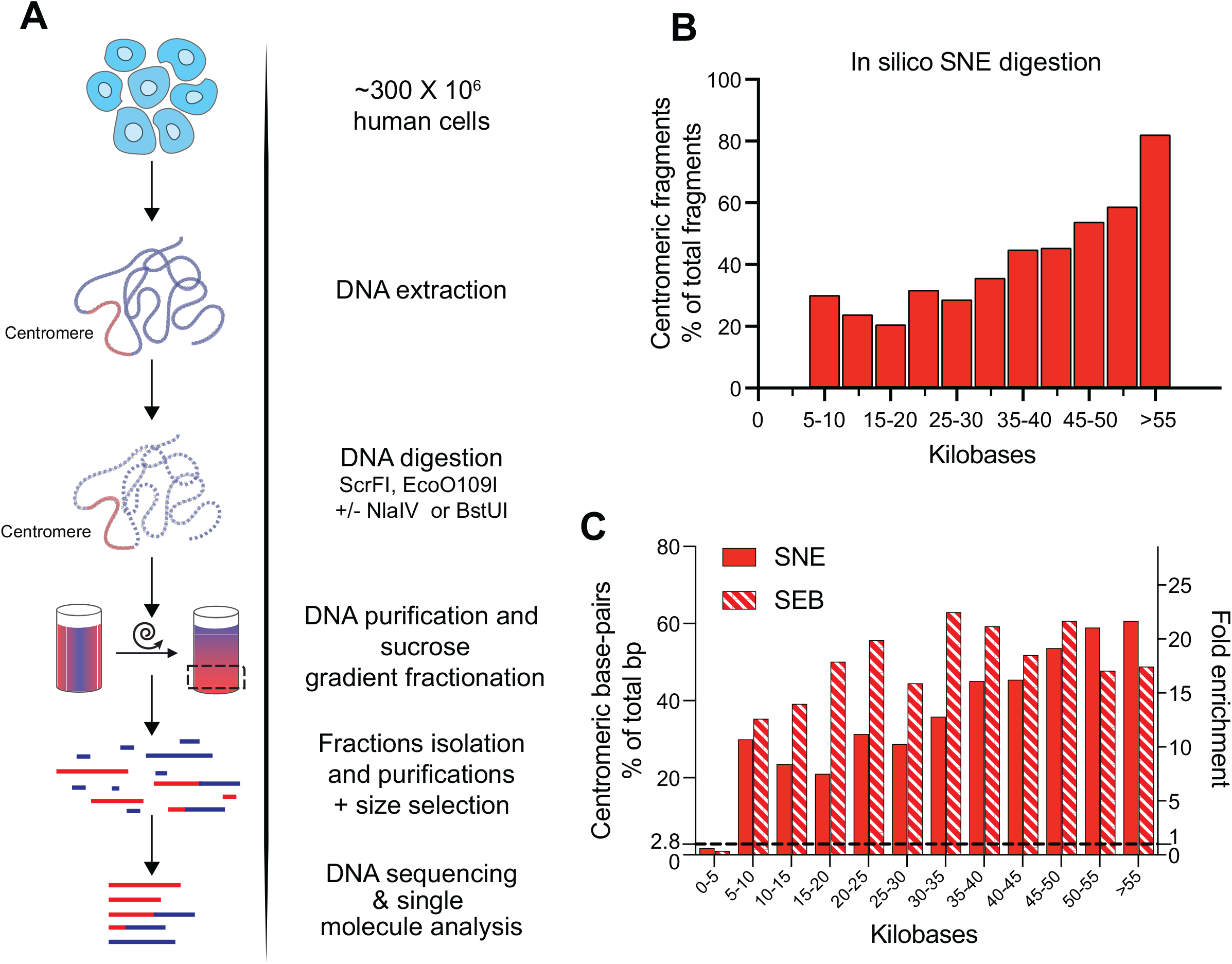
A restriction enzyme-based method to enrich and purify centromeric DNA from human cells. **A**. Schematic representation of the experimental design. **B**. Predicted distribution of the percentage of centromeric fragments in the indicated size bins after *in silico* digestion of the reference T2T-CHM13v1.0 genome with the SNE enzyme combination. Y-axis represents the percentage of centromeric fragments over total fragments in each length range. **C**. Distribution of centromeric base-pairs according to predicted fragment length after *in silico* digestion of the reference T2T-CHM13v1.0 genome with the SNE or SEB enzyme combinations. The y-axis on the left represents the percentage of centromeric base-pairs over total base-pairs in each length range. The dotted line at 2.8 % represents the percentage of centromeric base-pairs in the reference genome, corresponding to the expected fraction of centromeric DNA in a theoretical non-enriched sample. The y-axis on the right reports the fold enrichment in centromeric base-pairs over the non-enriched sample (∼2.8% of centromeric base-pairs in the reference genome).

## RESULTS

### *In silico* digestion of a human reference genome

Taking advantage of recent progress in the determination of the sequence of human centromeres, we performed *in silico* digestions of the T2T-CHM13v1.0 reference genome (Nurk *et al*, 2021) using restriction sites from a panel of 240 commercially available restriction enzymes. We then verified the size distribution of fragments deriving either from centromeric or non-centromeric regions. Based on this analysis we identified 2 candidate enzyme combinations (ScrFI + NlaIV + EcoO109I and ScrFI + EcoO109I + BstUI, hereafter named SNE and SEB respectively) that are predicted to cut non-centromeric DNA at high rate while digesting the centromeric regions at low frequency (**Figure S1A, B**). In both combinations, about half of centromeric DNA is digested into lower molecular weight (LMW) fragments (**Figure S1A, B**). However, with the SNE combination a high level of enrichment in centromeric DNA fragments is predicted in the HMW range (up to >80% of centromeric fragments >55 kb) (**Figure 1B**) corresponding to an enrichment in base-pairs up to 60% (**Figure 1C**). The SEB combination also showed a high percentage of centromeric fragments (40 to 60%) and centromeric base-pairs, but more homogeneously distributed in the range >15 kb (**Figures 1C** and **S1C**). Considering that in the reference genome the centromere content is about 2.8%, both combinations reach a fold enrichment in base-pairs of > 20-fold.

### Centromeric DNA purification from human cells

To test these predictions, we extracted and digested DNA from a pseudo-diploid, colorectal cancer cell line (DLD-1) with the SNE enzyme combination. The digested DNA underwent size fractionation by sucrose-gradient ultracentrifugation (20% to 40% sucrose weight/volume) and the collected fractions were used for dot-blot hybridization with a centromeric probe (CENP-B box) (**Figure 2A**). As a control we used a probe targeting the Alu repeats, an element which is widespread across the genome and not disproportionately abundant at centromeres. Indeed, short (250-300 bp) Alu sequences occupy about 307,000 kb of the genome (∼11%) (Deininger, 2011; Hoyt *et al*, 2021), but less than 20 elements per Mb are present at centromeres, only within divergent alpha satellite (Altemose *et al*, 2021a). Compared to unfractionated genomic DNA (gDNA), fractions 3 and 4 show the highest level of enrichment in centromeric DNA (about 20-fold), while Alu repeats were homogenously distributed among fractions (**Figure 2B, C**). The abundance of centromeric sequences in these fractions was also confirmed by qPCR for both the SNE and the SEB combinations (**Figure S2A**), further proving that the candidate enzyme mixes can be combined with size fractionation to enrich in centromeric DNA.

**Figure 2.**
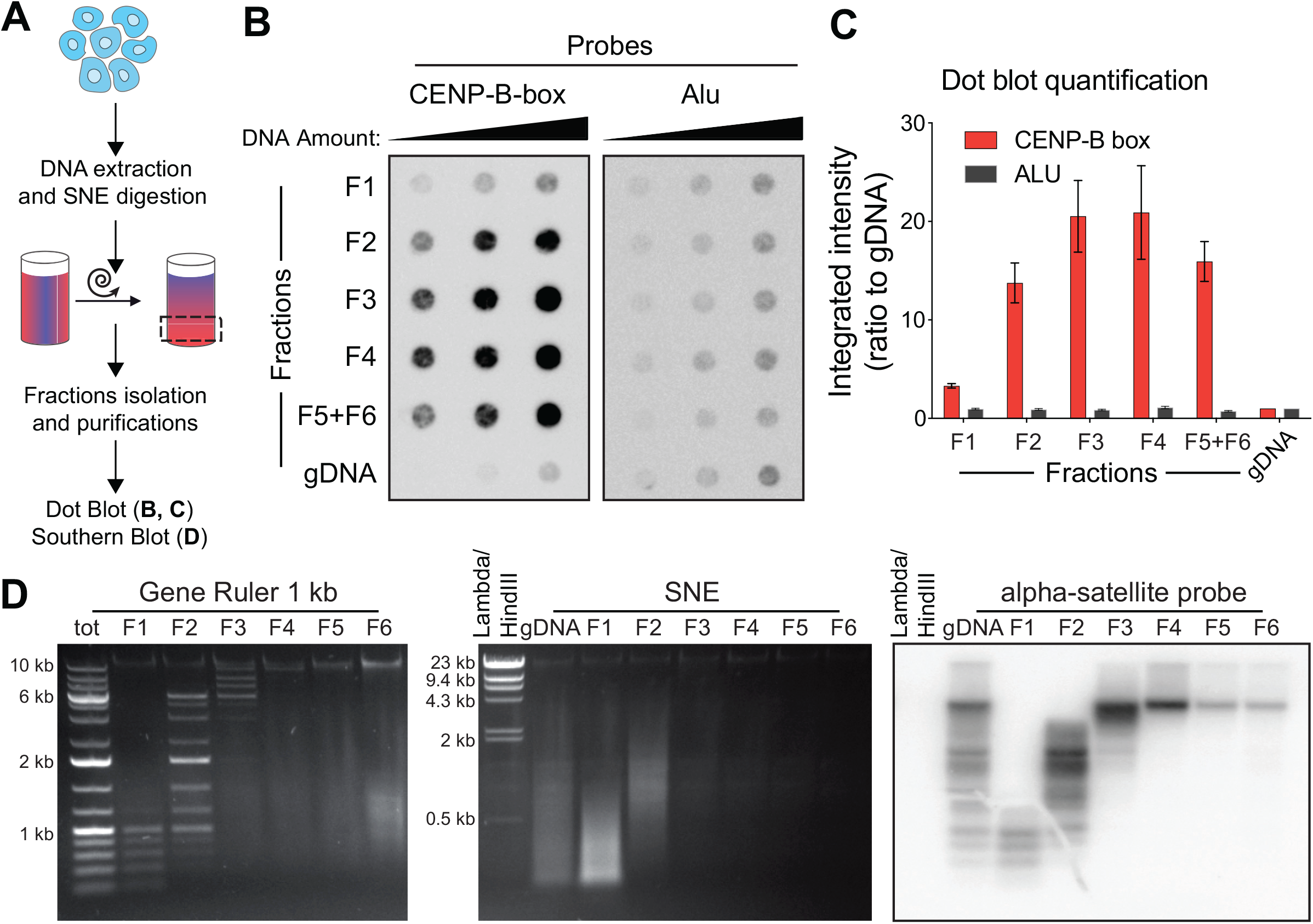
Centromeric DNA is enriched in the high-molecular weight fractions. **A**. Schematic representation of the experimental design. **B**. Dot-blot detecting the abundance of centromeric DNA (measured by signal intensity with a CENP-B box DNA probe, left membrane) in different sucrose gradient fractions (F1 to F4; F5+F6 is a pool of fractions F5 and F6) and in unfractionated genomic DNA (gDNA). A specific probe for the Alu repeat was used as a control (right membrane). In both membranes increasing amounts of DNA were loaded (50, 100 and 200 ng). **C**. Quantification of the dot-blot showed in B; signal is reported as a ratio to gDNA. The average for the different amounts of DNA is reported. Error bars represent the standard error of the three DNA amounts. **D**. Left: agarose gel electrophoresis performed on a molecular weight marker (Gene Ruler 1 kb), separated in the sucrose gradient showing efficient size separation; “tot” represents the unfractionated marker and F1 to F6 represent the different fractions. Middle and right: agarose gel electrophoresis of the sucrose fractions of a genomic DNA sample digested with the SNE combination and corresponding Southern blot after hybridization with an alpha satellite probe. “gDNA” represents the digested unfractionated sample and F1 to F6 represent different fractions. Lambda DNA digested with HindIII was also used as size control.

Restriction-based enrichment methods have been successfully used for telomeres since telomeric repeats do not contain restriction sites (including the recognition sites of our selected enzymes), therefore carryover of telomeric DNA may result in a decrease in the desired enrichment in centromeric DNA. To verify whether the centromere-enriched fraction also contained high amounts of telomeric DNA, we digested another batch of DLD-1 genomic DNA with the SNE enzyme combination, with the addition of a purified enzyme capable of cutting within telomeric repeats (T-EN) (Yoshitake *et al*, 2010). Following hybridization with centromeric or telomeric probes, we observed that while telomeric DNA is also detected mostly in the fractions F3 and F4, centromeric DNA still appears to be dominant (as expected due to its abundance over telomeric DNA). Addition of T-EN successfully depletes most of the telomeric signal (**Figure S2B, C**), making this approach also feasible in cell lines characterized by very long telomeres (e.g. ALT cell lines as U2OS).

To obtain information on the size distribution of the fragments resulting from digestion and fractionation, genomic DNA from diploid, non-transformed human hTERT RPE-1 cells was digested with SNE or SEB and analyzed by Southern blot using an alpha-satellite specific probe (**Figure 2D** and **S2D**). While the bulk of digested DNA is in fraction F1 and F2 (visualized as a smear in the agarose gel) and almost invisible in the HMW fractions (F3-F6), the centromeric signal is detected mostly in fractions F3 and F4 (**Figure 2D**). Although this type of gel does not allow high resolution in the HMW range, fractions F4-F6 appear to be > 10 kb long, which makes them suitable for approaches requiring long uninterrupted DNA molecules, such as long read sequencing or direct visualization with electron microscopy. As predicted *in silico*, centromeric DNA is also detected at LMW (F1 and F2), indicating that about half of centromeric DNA is digested into shorter fragments. Similar results were obtained for the SEB combination (**Figure S2D**). Hybridization with a telomeric probe shows that telomeric DNA is collected mostly in F2 and HMW fractions (F3-F6) are nearly devoid of telomeres while being rich in centromeric DNA (**Figure S2E**).

### Assessment of centromeric DNA enrichment by sequencing and single molecule visualization

Next, for both the SNE and SEB digestions, the two fractions appearing more enriched in centromeric DNA (F3 and F4) were sequenced with an Illumina NovaSeq 6000 system together with a low-enrichment fraction (F2) and an undigested, unfractionated DNA sample as control (hereafter referred as WGS) (**Figure 3A**). The resulting reads were then mapped on the T2T genome [T2T-CHM13v1.0 (Nurk *et al*, 2021)]. The reads were counted as centromeric when aligning within the genomic coordinates (**Table S1**), that contain both homogeneous HORs and monomeric/divergent alpha satellites, hereafter defined together as “centromeric regions”. In the SNE combination, about 54% of the reads map on centromeric regions in both F3 and F4 fractions, while only ∼3% of F2 and WGS reads are centromeric (**Figure 3B**). This corresponds to an approximately 18-fold enrichment in centromeric DNA compared to WGS. Similarly, we detect over 40% of centromeric reads in both in F3 and F4 for the SEB combination (**Figure S3A**).

**Figure 3:**
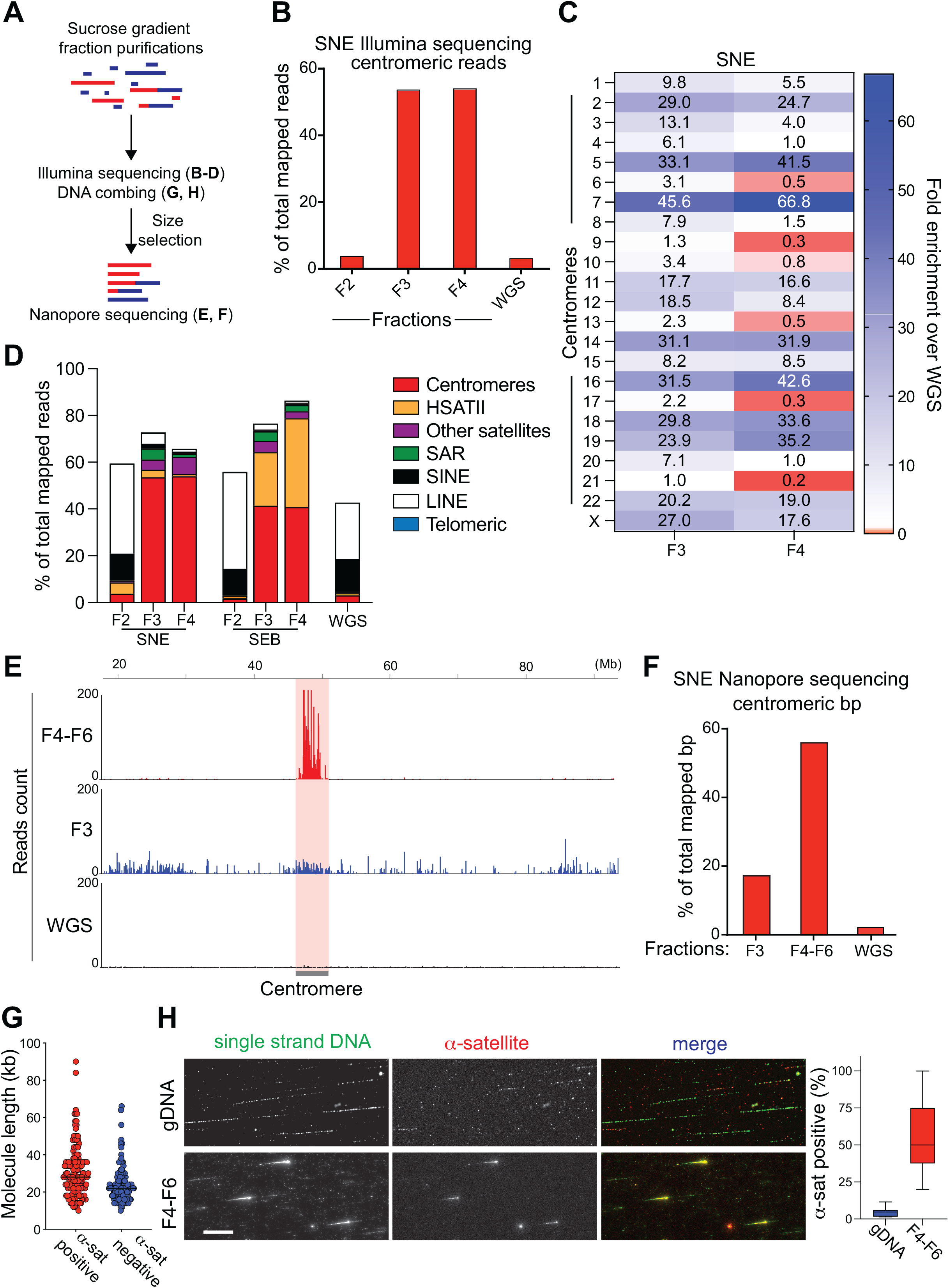
Enriched centromeric DNA is suitable for short or long-reads sequencing and single molecule analysis. **A**. Schematic representation of the experimental design **B**. Quantification of Illumina reads mapping in centromeric regions (as defined in Table S1), after SNE digestion and sucrose gradient separation (F2, F3 and F4) and in an undigested unfractionated sample (WGS). Read counts are reported as a percentage of total mapped reads. **C**. Enrichment in centromere-derived reads after Illumina sequencing across the different centromeres in fractions F3 and F4 after SNE digestion. Enrichment is expressed as a ratio to the read counts in the WGS sample. **D**. Quantification of Illumina reads mapping on different repeat families in fractions F2, F3, F4 after digestion with SNE or SEB enzyme combinations and in an undigested unfractionated sample (WGS). Read counts are reported as a percentage of total mapped reads. **E**. Coverage profiles of the centromeric region of chromosome 5 after Nanopore sequencing of an undigested sample (WGS), fraction 3 and a pool of fractions F4,5,6 (F4-F6). Genomic coordinates (Mb) are reported on top. **F**. Quantification of base-pairs from Nanopore reads that map within the centromeric regions (as defined in Table S1) after SNE digestion and sucrose gradient (fraction F3 and in a pool of fractions F4, F5, F6) and in an undigested unfractionated sample (WGS). Base-pair counts are expressed as percentage of total number of mapped base-pairs. **G**. Graph shows the size distribution in kb of DNA fibers positive or negative to a α-satellite probe. Each dot is a DNA fiber. Small fibers of less than 10 kb are not detected by microscopy. **H**. (Left) Representative images of DNA fibers hybridized with a single strand antibody and α-satellite probe in the indicated condition. Scale bar is 20 μm. (Right) Bar graph shows the averaged percentage of α-satellite-positive fibers in the enriched sample (F4-F6 fraction) or unfractionated DNA (gDNA) from different fibers (N = > 130). Error bars show standard deviation.

To verify efficiency of the restriction digestion, the Illumina sequencing data were tested for the presence of restriction sites within the reads, indicative of an incomplete digestion. For the F3 and F4 fractions of the SNE combination we detected very low level of intact restriction sites (<5% of total sites observed by WGS), indicative of a near-complete digestion efficiency (**Figure S3B**). Therefore, although the ScrFI and NlaIV enzymes are CpG DNA methylation-sensitive, their digestion rate seems largely unaffected, suggesting that most of these sites are unmethylated. In the SEB combination we detected a slightly higher fraction of undigested sites for the BstUI enzyme (15% and 10% in F3 and F4, respectively) (**Figure S3B**), possibly due to the increased effect of DNA methylation protection for this restriction site that contains two CpG dinucleotides.

To avoid the influence of potential mapping artifacts, we performed a k-mer based analysis aimed at identifying the reads containing alpha satellite sequence, while not relying on alignment to a reference assembly (see Methods). 45-50% of reads were identified as alpha satellite in the SNE sample (**Figure S3C**), a value that is compatible with the ∼54% of reads mapping within centromeric regions, where not all DNA is alpha satellite [for example, transposable elements (TEs) are present within arrays of divergent alpha repeats at a frequency of > 90 TEs per Mb (Altemose *et al*, 2021a)].

Centromeres of different chromosomes are characterized by different HORs on which reads can be differentially mapped thanks to the recent improvement in the assembly of human centromeres (Logsdon *et al*, 2021; Miga *et al*, 2020; Nurk *et al*, 2021). Therefore, we verified if centromere-derived reads in the enriched fractions are homogenously distributed across chromosomes or if some centromeres are more represented than others, considering that a large fraction of centromeric DNA is digested to LMW and therefore not further analyzed. Our analysis on the SNE digestion reveals that the distribution of the centromeric reads is heterogeneous, with some centromeres being largely overrepresented (e.g.: ∼67-fold enrichment for centromere 7) compared to the undigested, not fractionated WGS (**Figure 3C**). Overall, 17 out of 23 centromeres are enriched by at least 5-fold in either F3 or F4, while only chromosomes 21 and 9 show no enrichment. Performing the same analysis on the SEB digestion also shows inter chromosomal heterogeneity, but with a different pattern of centromeric reads distribution compared to SNE (**Figure S3D**).

The mapped reads were then further analyzed to test for the presence of other repetitive DNA families using RepeatMaskerV2 annotations: fractions F3-F4 of the SNE digestion led to about 10-12% of reads mapping on other satellite DNA, notably belonging to the families of satellite II (“HSatII”) and SAR (more recently recategorized as “HSat1A”) (Altemose *et al*, 2021a) (**Figure 3D**). Interestingly, the same fractions of the SEB digestion mix show a much higher abundance of the pericentric HSatII (23% and 38% for F3 and F4, respectively, with a fold enrichment up to 41-fold compared to WGS), with only a minor decrease in the fraction of centromeric DNA (**Figure 3D**). As expected, short interspersed mobile elements like SINEs and LINEs are underrepresented in the high molecular weight fractions and tend to remain in the F2 both for SNE and SEB (**Figure 3D**). Only very low levels of telomeric DNA were identified, mainly in F2, as expected from RPE-1 cells and from the Southern blot results (**Figure S2E**).

Since long reads sequencing techniques have become crucial for the dissection of repetitive arrays like centromeres, we tested the applicability of the centromere enrichment method to Nanopore sequencing, with particular interest in the centromere enrichment level and in the read length that can be obtained (**Figure 3A**). Following DNA digestion with SNE and size fractionation, sucrose gradients fractions F4 to F6 (F4-F6) were pooled and sequenced with the Oxford Nanopore Technologies system. In parallel, fraction F3 and an undigested sample (WGS) from the same cell line were also sequenced. A capillary electrophoresis analysis showed that while the mass of F4-F6 consists mainly of >50 kb fragments, contamination with molecules down to ∼1 kb is also present (**Figure S3E**, red line), which negatively affected the average read length in the output of Nanopore sequencing (data not shown). Therefore, prior to sequencing we used a size selective precipitation method (see Material and Methods) to efficiently remove <10 kb DNA molecules and to further enrich the sample in long DNA fragments (**Figure S3E**, blue line). Sequencing of this sample led to a N50 of ∼22 kb (50% of the sequenced base-pairs are within reads >22 kb long) with about 40% of reads longer than 15 kb (**Figure S3F**). Following this additional step of size purification, at most centromeric regions there is a strikingly higher coverage in F4-F6 fractions compared to WGS (**Figure 3E**). Specifically, F4-F6 shows a >26-fold enrichment in overall centromeric DNA compared to WGS, with about 55% of the total sequenced base-pairs being of centromeric origin (**Figure 3F**). F3 has an enrichment level of ∼8-fold, with 17.5% of the base-pairs deriving from centromeric regions (**Figure 3E, F**). In summary, the restriction digestion-based centromere selection method can efficiently be used in combination with Nanopore sequencing to reach unprecedented levels of enrichment in centromeric DNA while also preserving several kb long reads at a fraction of the cost of WGS.

We finally tested the feasibility of single-molecule direct visualization on the centromere-enriched sample by microscopy. To this end, DNA fibers from the pool of fractions F4 to F6 after SNE digestion and undigested unfractionated samples (gDNA) were subjected to DNA combing assay coupled with a mix of fluorescent probes against α-sat DNA (**Figure 3A**), as previously done (Giunta *et al*, 2021). Here we observed that in the pooled F4-F6 sample the DNA fibers have a mean length of ∼28 kb, with the one positive for the α-sat probe being slightly longer (median distribution of centromeric vs non-centromeric of 28 kb vs 22 kb, mean ∼34 kb vs ∼23 kb, respectively) (**Figure 3G**). In agreement with our sequencing data, in the F4-F6 sample, DNA fibers > 30 kb consist of about ∼56 % of alpha satellite array while in the gDNA sample is ∼4 %, as expected (**Figure 3H**). This result indicates our enrichment protocol is suitable for visualization and analysis of single DNA molecules.

## DISCUSSION

In this manuscript, we provide a simple and reliable method to enrich for centromeric DNA independently from the binding of proteins (unlike ChIP/Cut&Run on centromeric proteins). We show that our approach is compatible with DNA sequencing (either short-reads and long-reads sequencers) and for direct visualization of single molecules.

This method provides several advantages. First, it has great potential to make sequencing of centromeric DNA more affordable and efficient. From our results we can estimate that, to obtain an average coverage across centromeres of 5X, WGS would require sequencing of over 15 Gb, while the enriched F4-F6 would only need about 0.6 Gb, with a striking decrease in sequencing cost and time. This new development has great possibilities of application and is particularly timely, as the study of centromeric DNA has just entered a new genomic era thanks to the fine mapping and assembly of their repeats (Altemose *et al*, 2021a; Logsdon *et al*, 2021; Miga *et al*, 2020; Nurk *et al*, 2021). However, we are just scratching the surface of the molecular characterization of centromeric DNA. Indeed, even if human centromeres have been well-characterized by the T2T consortium, the data derive only from on one cell line, while centromeres can vary across individuals and can be drastically altered (e.g. length, epigenetic status, organization) in pathological conditions associated with genome instability, such as cancer. We therefore envision that the study of centromeric DNA will be in high demand in the near future, and our method can provide a valuable tool to facilitate centromere sequencing.

Second, our data revealed heterogeneity in the level of enrichment obtained among different centromeres (**Figures 3C** and **S3D**). This observation highlights that different enzyme combinations or choice of fractions can be used to focus the enrichment on selected centromeres. Since population diversity in the size and sequence composition of centromere has been described (Miga, 2019), it is possible that the observed enrichment distribution may vary depending on the cell line used. Furthermore, this approach can also be applied to study pericentromeric regions, both for sequencing and direct DNA visualization. For example, for studies focusing both on pericentromeres and centromeres, the SEB combination has the potential to obtain an enriched sample where almost 80% of the DNA consists of HSatII and alpha satellite (**Figure 3D**).

While preparing this manuscript, a similar restriction-digestion method combined with agarose gel separation to enrich for centromeric DNA was presented in a pre-print report (Altemose *et al*, 2021b). Here the authors used a different enzyme combination (MscI and AseI, hereafter referred as MA) from the one presented here. *In silico* digestion with MA revealed that centromeric DNA is better preserved compared to SNE or SEB combination (**Figure S4A**), but overall the percentage of centromeric fragments is lower compared to our enzyme combinations (**Figures 1B, S1C** and **S4B**). A likely explanation of this result is that non-centromeric DNA is less fragmented with the MA combination compared to SNE or SEB, leading to a higher contamination of non-centromeric fragments at HMW fractions. Considering all fragments > 10 kb the percentage of centromeric fragments in SNE and in MA is lower than SEB, but when the cutoff is set at >25 kb SNE is the best performing enzyme combination with an increasing prevalence of centromeric fragments in longer fragment ranges (**Figures 1B, S1C** and **S4B**). On this regard, the sucrose gradient isolation that we performed here allowed isolation of fragments above 10-15 kb, but, according to our *in silico* prediction, it is possible to have an even purer sample if we can obtain a higher cutoff. While the *in silico* prediction can vary significantly from what really observed in cells, the MA enzymes combination represents a valid alternative to the one presented here when preservation of total centromeric DNA, but not its purity, is the main target. It is important to note that all three enzyme combinations show inter-chromosomal heterogeneity in centromeric fragments prevalence across centromeres, with SNE being the most heterogeneous and SEB the least variable (coefficients of variation of 111%, 81% and 76% for SNE, MA and SEB respectively, measured taking into account the abundance of all fragments > 20 kb) (**Figure S4D**).

Finally, our method to enrich and purify human centromeres is suitable for direct visualization of single DNA molecules. This includes analysis of replicating DNA fibers aimed to study replication fork dynamics and stability using techniques as DNA combing (**Figure 3G, H**). Such approaches that rely on the usage of DNA probes to label specific regions like centromeres, despite being feasible (Giunta *et al*, 2021; Li *et al*, 2018; Mendez-Bermudez *et al*, 2018), may pose technical issues. By having more than half of the DNA fibers of centromeric origin (**Figure 3H**), it is possible to bypass the usage of specific labeling or, depending on the enzyme combination, perform replication studies on specific centromeres. This is even more important in techniques in which the usage of fluorescent probes is not feasible as EM or atomic force microscopy. For example, following *in vivo* psoralen crosslinking, the enriched centromeric DNA can be further processed for EM to study the topological structures and replication intermediates at centromeres, as done for telomeres (Mazzucco *et al*, 2020; Griffith *et al*, 1999). This has the potential to shed light on the architecture and replication intermediates that are present at centromeric regions and to understand how centromeric DNA binding proteins might modulate their topology and structure.

In conclusion, our method represents an invaluable tool for the study of human centromeric repeat arrays.

## MATERIAL AND METHODS

### Cell lines

All cells were maintained at 37°C in a 5% CO_2_ atmosphere. Immortalized hTERT RPE-1 cells were cultured using DMEM:F12 medium containing 10% Fetal Bovine Serum (BioSera), 0.123% sodium bicarbonate, and 2 mM L-glutamine. DLD-1 cells were grown in DMEM medium containing 10% Fetal Bovine Serum (BioSera).

### Purification of telomere-digesting chimeric endonuclease (T-EN)

The telomere-digesting TRAS1EN-TRF1 chimeric endonuclease (T-EN) was expressed from a pET21b plasmid kindly provided by H. Fujiwara (University of Tokyo) (Yoshitake *et al*, 2010). Briefly, histidine-tagged T-EN was expressed in BL21-CodonPlus-RIL competent cells at 20°C, and purified by affinity chromatography on a 5 ml His-Trap FF crude column (GE Healthcare), the protein was further purified by gel filtration using a HiLoad Superdex 200 16/600 column (GE Healthcare).

### DNA extraction and digestion

Genomic DNA was extracted from 300-400 million cells as described in (Mazzucco *et al*, 2020) Briefly:

1. Cells were trypsinized, washed twice in PBS 1X and resuspended in TNE buffer (10 mM Tris-HCl, 1mM EDTA pH8, 10 mM NaCl).
2. Cells were lysed by adding one volume of TNES buffer (TNE + 1% SDS) supplemented with RNaseA (Invitrogen cat #12091021) at final concentration of 100 μg/mL, and incubated at 37° for 30 minutes.
3. Proteinase K treatment (Invitrogen cat #25530049) was performed overnight at 37° at a final concentration of 100 μg/ml.
4. DNA was extracted with one volume of Phenol:Chloroform:Isoamylalcol (25:24:1) (Sigma Aldrich cat#77617). After centrifugation at 3500 g for 5 minutes, one volume of chloroform was added to the aqueous phase.
5. After centrifugation at 3500 g for 5 minutes, the DNA in the aqueous phase was precipitated with 0.1 volume of sodium acetate 3M pH 5.2 and one volume of Isopropanol.
6. After washing with 70% ethanol, DNA was gently resuspended in 1 ml of Tris-HCl 10 mM pH 8.0.
7. 2.5 mg of DNA were resuspended in 20 mL of 1X CutSmart Buffer (NEB cat#B7204S) and incubated at RT for one hour on a rotating wheel.
8. Digestion was carried out over night at 37° using 400 units each of ScrFI and EcoO109I and with 400 units of NlaIV or BstUI (New England Biolabs). When applicable, 1 μM of T-EN enzyme (telomere digesting) was added to the digestion mix.
9. Digestion products were purified with one step of phenol:chloroform:isoamylalcol (25:24:1) purification and precipitated with isopropanol and sodium acetate, as above. DNA was resuspended in 4.5 mL of TE 1X.

### Sucrose gradient fractionation

1. Sucrose gradients were prepared with 8 ml each of 40%, 30% and 20% sucrose solutions in TNE buffer, carefully deposited sequentially on top of each other in Thickwall, Ultra-Clear tubes (Beckman Coulter cat #344058) compatible with SW32Ti rotor.
2. The digested DNA sample was split in 4 aliquots, each in a volume of 1.5 ml, and incubated at 50° for 5 minutes prior to loading each aliquot on a separate sucrose gradient.
3. The gradients were centrifuged at 4° in a SW32Ti rotor at 30100 rpm for 16 hours.
4. The fractions were collected as follows: the top 5.5 ml were collected as fraction 1 (F1) while the remaining F2 to F6 consisted of 4 ml each.
5. Fractions were concentrated using Amicon Ultra 15 ml centrifugal filters (MWCO=30 kDa, Merck, cat# UFC903024) performing 5-6 washes of the filter with Tris-HCl 10 mM pH 8.0. The sample (0.5-1 ml) was transferred to Amicon Ultra 0.5 ml Centrifugal Filters (MWCO=30 kDa, Merck, cat# UFC503096) and further concentrated to a final volume of 200 μl.

### qPCR, dot blot and Southern Blot

qPCR was performed using the LightCycler 480 (Roche) system with previously described primer pairs specific for alpha satellite DNA, as target (5’-TCCAACGAAGGCCACAAGA-3’ and 5’-TCATTCCCACAAACTGCGTTG-3’) and for the 18S rDNA, as reference (5′-CTCAACACGGGAAACCTCAC-3 and 5′-CGCTCCACCAACTAAGAACG-3′). Fold enrichment was calculated with the ΔΔCt method as enrichment of the target sequence over the reference. For the dot blot experiments, 50, 100 and 200 ng of DNA from each fraction and from unfractionated genomic DNA were blotted on a membrane (Amersham Hybond -N+, GE Healthcare) using a BioDot apparatus (Bio-rad). Membranes were hybridized overnight at 42°C with digoxigenin-3’-labeled oligos as probes specific for CENP-B boxes (5’-ATTCGTTGGAAACGGGA -3’), Alu repeats (5’-ATACAAAAATTAGCCGGGCG -3’) or telomeres (5’-TAACCCTAACCCTAACCCTAACCCTAA -3’). Signal detection was performed with CDP Star solution (Roche) and imaged with a Chemidoc imaging system (Biorad).

For Southern blot analysis, 1:1000 of each fraction together with 300 ng of unfractionated, digested gDNA were loaded on a 0.8% agarose gel in 0.5X TBE. Electrophoresis was performed at 5 V/cm for 90 minutes. After depurination, denaturation and neutralization, the DNA was blotted by capillarity on an Amersham Hybond-X (GE healthcare) membrane and crosslinked in a UV Stratalinker 1800 (Stratagene) with 1200 J of 254 nm UV. The membrane was pre-hybridized 1 hour at 65° in Church mix (500 mM NaPi pH 7.2, 1 mM EDTA pH 8.0, 7% SDS, 1% BSA). Hybridization occurred overnight in Church mix with a telomeric TTAGGG probe (Mazzucco *et al*, 2020) or centromeric probe (produced as described below). After three washes in Church wash buffer (40 mM NaPi pH 7.2, 1 mM EDTA pH 8.0, 1% SDS), radioactive signal was impressed on a FUJIFILM Storage Phosphor screen for 5 hours and acquired with Typhon Trio (GE healthcare).

Centromeric probe for Southern was produced by apha-^32^P-dCTP-labelling (Prime-a-Gene Labeling System, Promega cat #U1100) of a ∼300 bp PCR product obtained with primers 5′-CAGAAACTTCTTTGTGATGTGTGC-3′ and 5’-GTTTTTATGGGAAGATATTTCCT-3’ on a template of human genomic DNA.

### Libraries preparation and sequencing

Illumina sequencing libraries were prepared from unselected genomic DNA (WGS) and from the same fractions F2, F3 and F4 that were analyzed by Southern blot. After shearing to an average fragment size of 300 bp with a Covaris ME220 Sonicator, libraries were prepared with Kapa Hyper Prep kit (Roche) according to the manufacturer’s instructions with 12 amplification cycles. Pair-end sequencing was carried out with an Illumina NovaSeq6000 instrument.

Nanopore sequencing was performed from fractions derived from an independent digestion and sucrose gradient experiment. Before preparation of libraries for Nanopore sequencing, fractions F4 to F6 were pooled and 9 μg of this DNA was treated with Short Read Eliminator kit (cutoff <25 kb, Circulomics cat# SKUSS-100-101-01) to further remove contamination from shorter DNA fragments. Libraries were prepared from this sample, from fraction F3 and from total genomic DNA (WGS) using the Ligation Sequencing kit (Oxford Nanopore Technology, cat# SQK-LSK109). For all samples, sequencing was performed on a Spot-ON Flow Cell (R9.4.1) on a MinION Mk1B device.

Libraries were quantified with Qubit dsDNA HS Assay Kit (Thermo Fisher) and checked by capillary electrophoresis with a TapeStation 4150 system (Agilent).

### Bioinformatic analysis

#### *In silico* digestion

The reference genome used is the T2T-CHM13v1.0, where the centromeric and non-centromeric regions were defined according to the ranges reported in Table S1. *In silico* digestion was performed by matching the occurrence of each restriction site sequence and replacing it with a line break. The lengths of the resulting strings were used to represent the size of digestion products. Distribution analysis and plotting was performed with RStudio (RStudio Team, 2019)

#### Illumina sequencing

Illumina reads from all the fractions and from WGS were downsampled to the same total read count. The estimate quantification of alpha-satellite-derived Illumina reads was performed by counting the reads containing at least two of the previously identified unique alpha 18-mers representative of the alpha satellite DNA variation in the human genome (Miga, 2017).

Illumina reads were mapped using bwa-mem algorithm of the BWA software package (Li, 2013; Li & Durbin, 2009) on the Telomere-to-Telomere T2T-CHM13v1.0 reference genome (Nurk *et al*, 2021). Reads mapping on centromeric regions were counted according to the ranges specified in the Table S1. Reads mapping on different families of repeats were counted according to the ranges defined by the track Repeat MaskerV2 retrieved by UCSC Table Browser (Karolchik *et al*, 2004) on the assembly T2T-CHM13v1.0.

#### Nanopore sequencing

Nanopore sequencing data was basecalled with Guppy version 4.0 with a high accuracy model (*dna_r9*.*4*.*1_450bps_hac*.*cfg*) and were mapped using Winnowmap 2.0 (Jain *et al*, 2020a, 2020b). Primary alignments were filtered using samtools (version 1.9, (Li *et al*, 2009)) option -F 2308. Samtools mpileup was used to quantify the number of mapped bases within centromeric regions (Table S1).

### DNA combing

Combing and FISH analysis was performed on genomic undigested DNA and on the pool of fractions F4 to F6 from the SNE digestion. DNA was diluted in 0,25 M MES buffer (pH 5.5) and the DNA/MES mix was combed onto silanized coverslips (Genomic Vision) using the Molecular Combing System (Genomic Vision). DNA fibers were denatured for 5 min in 1 N NaOH, followed by PBS (4 °C) wash and dehydration in increasing concentrations of ethanol (75, 85, and 100 %). Slides were hybridized overnight at 37 °C with a biotinylated RNA alpha-satellite probe [see (Giunta *et al*, 2021)] and washed 3 times with 50% formamide solution at RT. After 3 washes in 2X SSC and a quick wash in PBS, slides were incubated for 1h in blocking solution (blocking reagent Roche, 11096176001) at 37°C. Centromere signal was detected by alternating layers of avidin FITC (1:100, 434411, Thermo) and rabbit anti-avidin biotin conjugated (1:50, 200-4697-0100, Rockland laboratories) antibodies. Single stranded DNA was detected with rabbit anti single-stranded DNA antibody (1:2, JP18731, Tecan/IBL international) and anti-rabbit CyTM3 (1:250, 711-165-152, Jackson Immuno Research). Fibers were mounted in ProLong Gold antifade reagent (P36935, Invitrogen) and acquisition was performed with an epifluorescence microscope (Upright LEICA DM6000).

### Data sharing

Illumina WGS data are available on the SRA database at the accession number SRX5973263. All other sequencing datasets will be available at the accession number PRJNA765341.

## Acknowledgements

The authors would like to thank Haruhiko Fujiwara (University of Tokyo, Japan) for sharing reagents and Raphael Margueron (I. Curie) for sharing equipment. Glennis Logsdon (University of Washington, US), C. Bartocci (I. Curie) and all other Fachinetti team members for helpful suggestions. High-throughput sequencing was performed by the ICGex NGS platform of the Institut Curie supported by the grants ANR-10-EQPX-03 (Equipex) and ANR-10-INBS-09-08 (France Génomique Consortium) from the Agence Nationale de la Recherche (“Investissements d’Avenir” program), by the ITMO-Cancer Aviesan (Plan Cancer III) and by the SiRIC-Curie program (SiRIC Grant INCa-DGOS-4654).

## Author contributions

R.G. performed the *in silico* and *in vitro* digestion, dot blot, qPCR, library preparation for sequencing and data analysis; G.M. performed sucrose gradient purifications, southern blots and dot blots; F.C. performed the *in vitro* protein purification for telomere digestion; L.V. performed sucrose gradient purification and alternative methods to enrich for HMW; T.W. performed DNA fiber analysis; J.P. helped in the in silico digestion; R.G., Y.D. and D.F. conceived the experimental design; D.F. directed the research and provide financial support; R.G. and D.F. made figures and wrote the manuscript. All authors contributed to manuscript editing.

## Supplementary figure legends

**Supplementary Figure 1.**
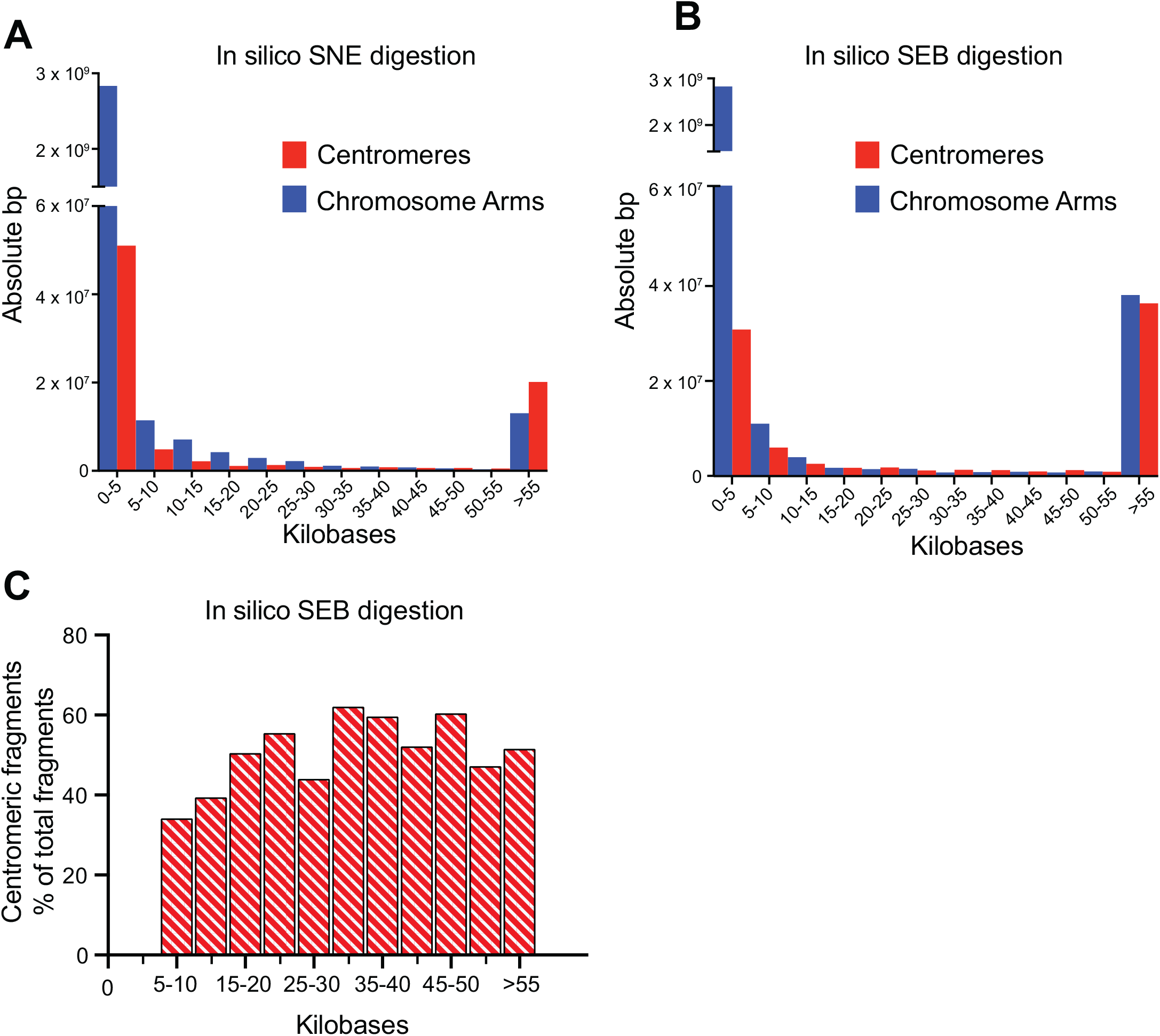
Distribution of centromere DNA after *in silico* digestion of a reference genome. Related to figure 1. **A-B**. Distribution of centromeric (red) or non-centromeric (blue) base-pair content of predicted fragments according to fragment length after *in silico* digestion of T2T-CHM13v1.0 genome with enzyme combinations SNE (A) and SEB (B). **C**. Distribution of predicted fragment lengths of centromeric fragments after *in silico* digestion of the reference T2T-CHM13v1.0 genome with the SEB enzyme combination. y-axis represents the percentage of centromeric fragments in each length range.

**Supplementary Figure 2.**
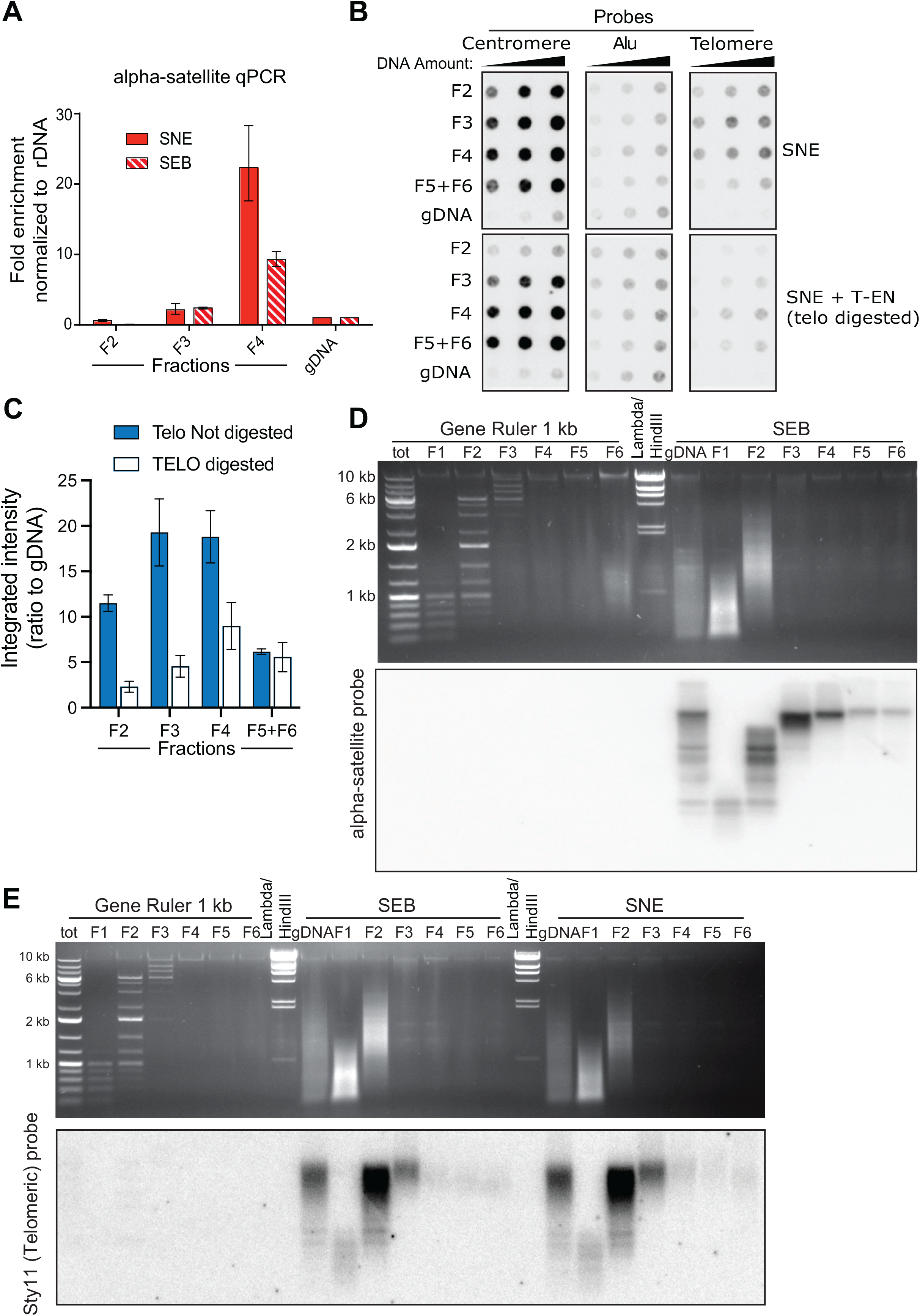
Centromeric DNA is enriched in the high-molecular weight fractions, which are deprived of telomeric DNA. Related to figure 2. **A**. qPCR analysis showing enrichment in centromeric DNA in the different sucrose fractions after digestion with SEB or SNE enzyme combination. Ct values were normalized to the signal from a ribosomal-DNA-specific primer pair. Fold enrichment is expressed over the undigested unfractionated genomic DNA sample. Error bars show standard deviation, n=3. **B**. Dot-blot to detect abundance of centromeric DNA (measured by signal intensity with a CENP-B box probe, left membranes) or telomeric DNA (right membranes) in different sucrose gradient fractions (F2 to F4; F5+F6 is a pool of fractions F5 and F6) and in unfractionated undigested genomic DNA (gDNA). A specific probe for the Alu repeat was used as a control (middle membranes). In all membranes increasing amounts of DNA were loaded (50, 100 and 200 ng). The top three membranes were loaded with samples digested with SNE combination enzymes (same as Figure 2B), while the bottom three membranes were loaded with samples digested with SNE + telomere specific endonuclease (T-EN). **C**. Quantification of the telomeric signal from the dot-blot showed in B; signal is reported as a ratio to gDNA. Bars represent the average of the different amounts of DNA. Error bars represent the standard error of the three DNA quantities. **D**. Agarose gel electrophoresis performed on genomic DNA digested with the SEB combination (top) and corresponding Southern blot after hybridization of the membrane with an α-satellite probe (bottom). “gDNA” represents the unfractionated sample and F1 to F6 represent different fractions. Efficient size separation is shown by the fractionation in sucrose gradient of a molecular weight marker (Gene Ruler 1 Kb). **E**. Agarose gel electrophoresis and corresponding Southern blots performed on genomic DNA digested with the SNE and SEB combinations, after hybridization with telomeric probe. “gDNA” represents the unfractionated sample and F1 to F6 represent different fractions. A molecular weight marker was used as a control and tested by agarose gel electrophoresis (Gene Ruler 1 Kb) proving the efficiency of sucrose gradient fractionation.

**Supplementary Figure 3:**
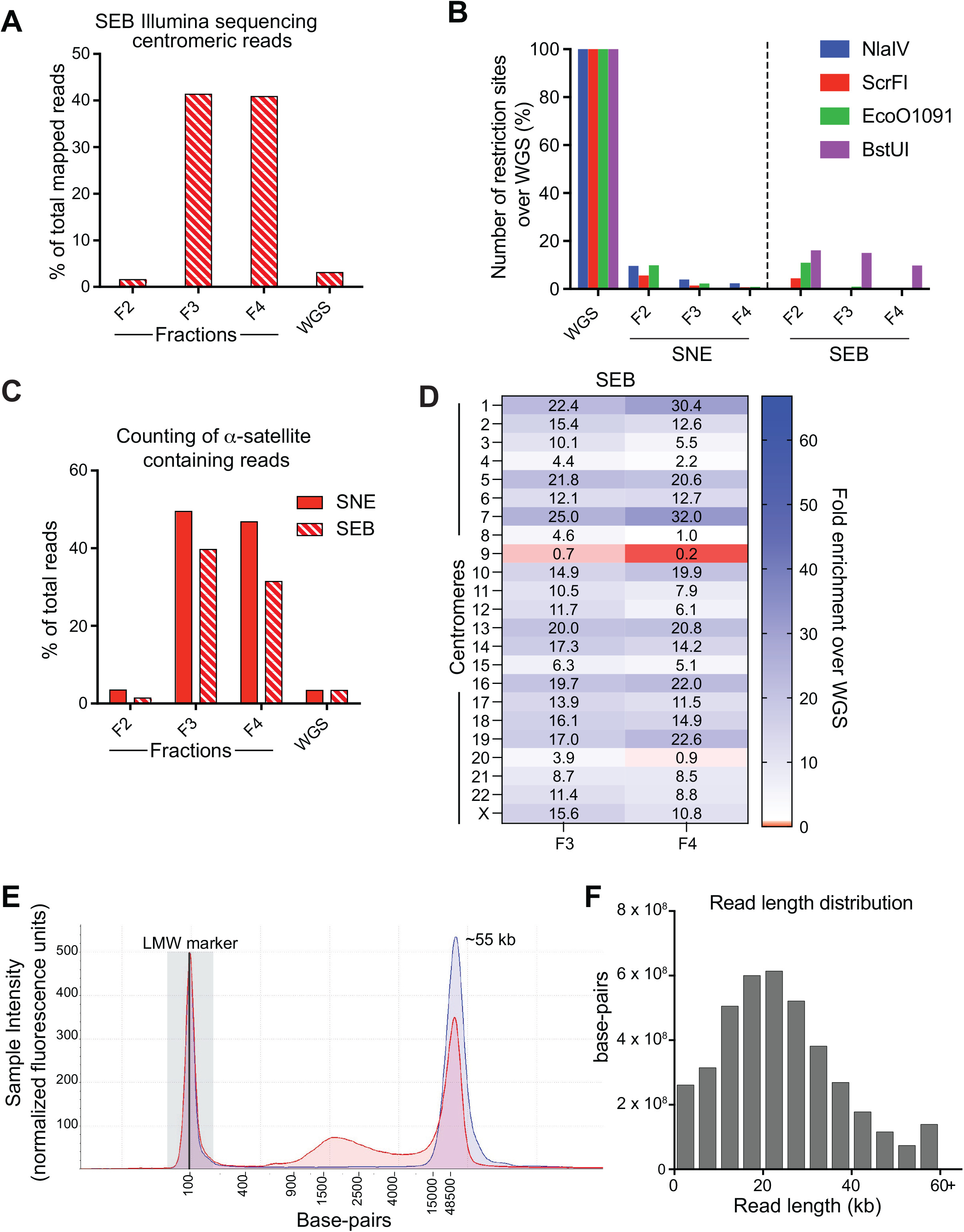
Enrichment in centromeric DNA detected by sequencing. Related to figure 3. **A**. Quantification of Illumina reads mapping in centromeric regions (as defined in Table S1), after SEB digestion and sucrose gradient separation (F2, F3 and F4) and in an undigested sample (WGS). Read counts are reported as a percentage of total mapped reads. **B**. Quantification of uncut restriction sites identified within Illumina reads after digestion with SNE or SEB enzyme combinations and fractionation (fractions F2, F3, F4). Values are reported as % of the sites identified in the reads from an undigested unfractionated sample (WGS). **C**. Quantification of Illumina reads containing alpha satellite 18-mers, after SNE or SEB digestion and sucrose gradient separation (F2, F3 and F4) and in an undigested sample (WGS). Read counts are reported as a percentage of total reads. **D**. Enrichment in centromere-derived reads after Illumina sequencing across the different centromeres in fractions F3 and F4 after SEB digestion. Enrichment is expressed as a ratio to the read counts in the WGS sample. **E**. TapeStation electropherogram profiles of SNE-digested DNA after sucrose gradient fractionation and pooling of fractions F4 to F6, before (red line) and after (blue line) additional size selection by precipitation with the Short Read Eliminator kit. The bulk of DNA is within a peak at ∼55 kb. The peak at 100 bp (labelled “lower MW marker” and marked with a grey rectangle) corresponds to a calibrator added for comparison of the two samples. **F**. Distribution of base-pair content of Nanopore reads according to read length.

**Supplementary Figure 4:**
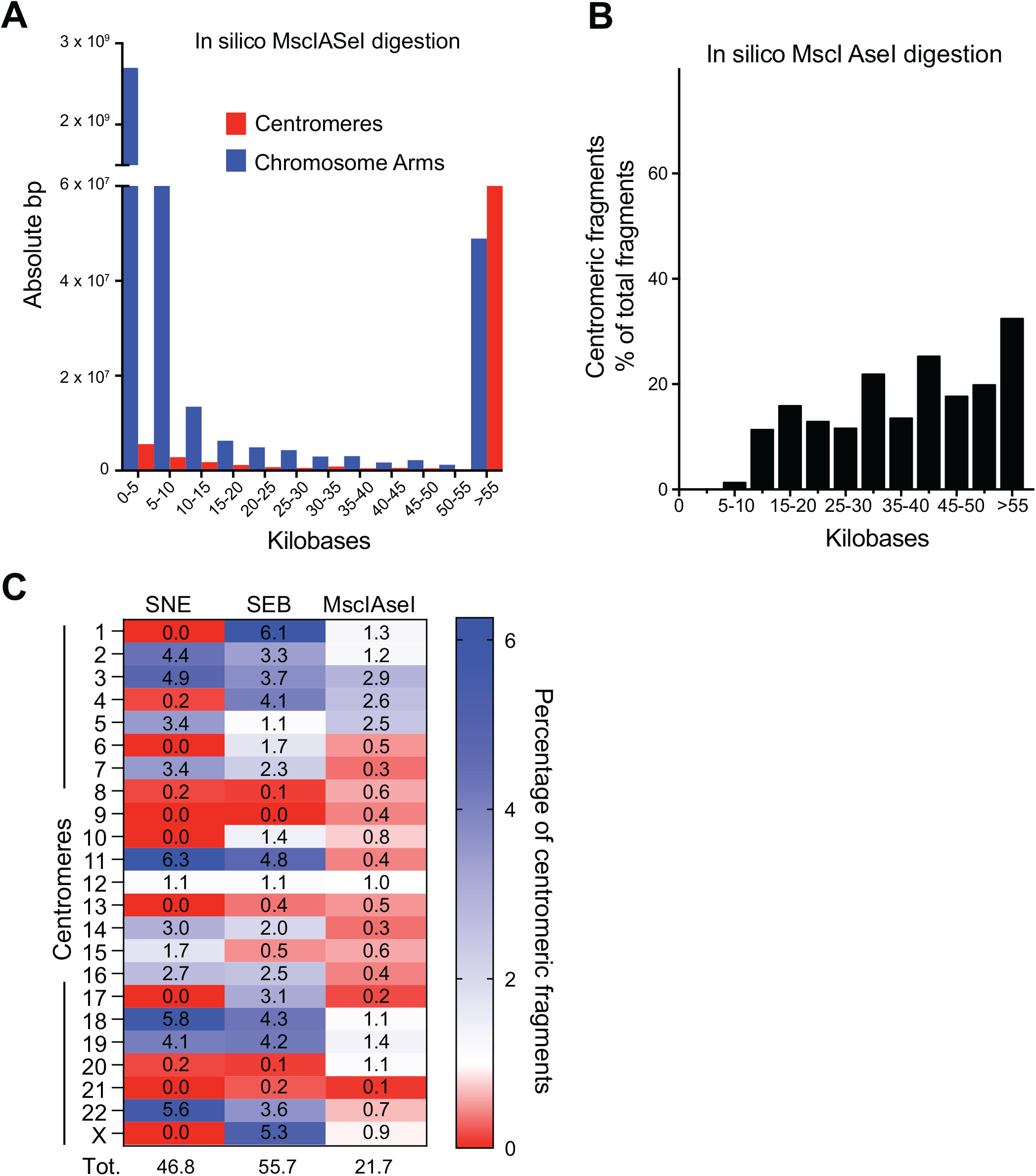
*In silico* digestion of a reference genome with an additional enzyme combination. Related to figure 1 and discussion. **A**. Distribution of centromeric (red) or non-centromeric (blue) base-pair content of predicted fragments according to fragment length after *in silico* digestion of T2T-CHM13v1.0 genome with MscI-AseI enzyme combination. **B**. Distribution of predicted centromeric fragment length after in silico digestion of the reference T2T-CHM13v1.0 genome with the MscI-AseI combination (black). y-axis represents the percentage of centromeric fragments in each length range. **C**. Distribution across different centromeres of the abundance of >20 kb centromeric fragments expressed as a percentage of all predicted fragments longer than 20 kb, after *in silico* digestion of the T2T-CHM13v1.0 genome with enzyme combinations SNE, SEB or MscI-AseI.

**Table S1:** Genomic coordinates on the T2T-CHM13v1.0 reference genome that define the boundaries of the centromeric regions.

| **Chromosome** | **start** | **end** |
| --- | --- | --- |
| chr1 | 121619339 | 126824297 |
| chr2 | 92297388 | 95077534 |
| chr3 | 90339864 | 96498631 |
| chr4 | 49705248 | 55303286 |
| chr5 | 46161723 | 50962189 |
| chr6 | 58243527 | 61413737 |
| chr7 | 60041897 | 64188432 |
| chr8 | 43846453 | 46920432 |
| chr9 | 44938599 | 47609552 |
| chr10 | 39068394 | 42061834 |
| chr11 | 50537301 | 54487964 |
| chr12 | 34485615 | 37369869 |
| chr13 | 16220369 | 18249463 |
| chr14 | 10125077 | 12841225 |
| chr15 | 15683284 | 18327600 |
| chr16 | 35743659 | 38213682 |
| chr17 | 23407665 | 27656858 |
| chr18 | 15542843 | 21125452 |
| chr19 | 24352654 | 30261056 |
| chr20 | 26281854 | 29953422 |
| chr21 | 11699867 | 12078398 |
| chr22 | 12326835 | 16903875 |
| chrX | 57593566 | 61247455 |

